# Extracellular cues accelerate neurogenesis of induced pluripotent stem cell derived neurons

**DOI:** 10.1101/2021.08.17.456634

**Authors:** Elizabeth R. Sharlow, Danielle C. Llaneza, Anna J. Mendelson, Garnett A. Mingledorff, Veronica M. Porterfield, Kathryn Salvati, Mark Beenhakker, George S. Bloom, John S. Lazo

## Abstract

Neurogenesis is a complex process encompassing neuronal progenitor cell expansion/proliferation and differentiation, followed by neuron maturation. *In vivo* models are most commonly used to study neurogenesis; however, human induced pluripotent stem cell-derived (iPSC) neurons are increasingly used to establish cellular models of human neurological processes. Unfortunately, the differentiation and maturation of iPSC-derived neurons varies in methodology, is asynchronous, and has restricted experimental utility because of extended differentiation/maturation times. To accelerate and standardize iPS neuronal maturation, we differentiated and matured feeder layer-free iPSC-derived neuronal cultures under physiological oxygen levels (5%), and modified the underlying extracellular matrix and medium composition. Our results demonstrate that calretinin gene expression occurred earlier under our optimized iPS conditions and the corresponding “neurogenesis burst” associated with proliferative expansion occurred more synchronously, reliably emerging two and three weeks after differentiation. As expected, the expression of mature neuronal markers (*i.e.*, NeuN+/Calbindin+) started at 4-weeks post-differentiation. qPCR microarray, western blot and single cell analyses using high content imaging indicated that 4-week iPS neuronal cultures were non-cycling with decreased expression of cyclin D1 and Ki67. Our data demonstrate that extracellular cues influence the kinetics of neurogenesis models and that feeder layer-free iPSC-derived neurogenesis can be reproducibly miniaturized.

## Introduction

Neurogenesis is the process by which new neurons are formed, or “born”, in the brain and it comprises neuronal progenitor cell expansion/proliferation and differentiation followed by neuron maturation. Prior to maturation, progenitor cells actively participate in the cell cycle wherein cells grow and divide; mature neurons no longer cycle. While neurogenesis is a critical component of development^1^, the process persists in some brain regions throughout the human lifespan^2^*(i.e.*, dentate gyrus of the hippocampus, olfactory lobe) as determined by BrdU incorporation, PCNA expression and the expression of neuronal maturation markers, such as doublecortin^3–5^.

Ongoing deliberations challenge the extent of neurogenesis in the adult human brain^6–7^. Several factors, including human tissue processing methods, may affect our ability to detect neurogenesis biomarkers^8^ and have therefore left the issue unresolved. Nonetheless, ample evidence from non-human systems (*i.e.*, rodents, non-human primates, songbirds)^9–14^ demonstrates that neurogenesis is not only detectable but can be enhanced by environmental cues such as exercise^15–16^, environmental enrichment (*i.e.*, learning)^17^ and engagement of the extracellular niche (*i.e.*, cell-cell interactions, extracellular matrix, ECM)^17^. Consequently, translating this “cuing” ability to promote healthy neurogenesis in neurological and/or neurodegenerative disorders has great appeal. Indeed, dysfunctional neurogenesis may be an early contributor to cognitive and behavioral decline in schizophrenia and Alzheimer’s Disease (AD)^18–21^. While *in vivo* models have been foundational in the study of neurogenesis, additional systems are needed to augment existing capabilities and enhance translation to patients.

Human induced pluripotent stem cell-derived (iPSC) neurons are increasingly being used as models to study neurological processes in healthy and disease states^22–23^. The utility of iPSC neurons stems mainly from their ability to recapitulate – at the cellular level – specific aspects of human neurodevelopment and neurodegenerative/neurological dysfunction previously only detected in patient-derived samples and/or non-human models. Hence, iPSC-derived neuron models, including organoids, “brain on a chip”, neurospheres, and various two/three dimensional systems are forming key components of neurophysiologically relevant systems to understand, for example, neurite outgrowth and neural network function^24–27^. Similarly, the diverse steps of neurogenesis (*i.e.*, neuronal progenitor cell (NPC) proliferation, migration and early differentiation of newly formed neurons) can be replicated by iPSC-derived neuron model systems^28^. Despite providing a physiologically relevant neurogenesis model, many studies using iPSC-derived neurons do not specifically evaluate neurogenesis and instead target differentiated or mature neurons to study later developmental processes such as synaptogenesis, electrophysiological signaling, or neurite outgrowth, or to study neurotransmitter receptor function^29–30^. The heightened focus on mature neurons may stem from suboptimal culturing conditions that often result in unsynchronized or unregulated proliferation of neuronal progenitor populations. Indeed, uncoordinated and unchecked proliferation complicates the study of neurogenesis in iPSC-based systems.

Herein, we aim to establish methods to refine and standardize neurogenesis in an astrocyte feeder layer-free human iPSC-based model system. Through the modification of extracellular cues (*i.e.*, underlying ECM, medium composition and culturing conditions (*i.e.*, 5% O_2_)), we accelerated the gene and protein expression of the post-mitotic protein, calretinin, providing a biomarker for human iPSC-derived neuron maturation. Using our high content imaging (HCI) platform and normal control NPCs, we have identified and quantified the “neurogenesis burst” associated with the rapid proliferation and early differentiation of normal iPSC-derived neuronal progenitor cell populations that lead to the development of mature, Calbindin+/NeuN+ iPSC neurons. This “neurogenesis burst” reproducibly occurs and coincides with the progressive expression of neuronal maturation markers. We were able to detect a subpopulation of mature iPSC-derived neurons (*i.e.*, Calbindin+/NeuN+) as early as 4 weeks post-differentiation, underscoring the capacity and robustness of our feeder layer-free culture system. Critically, our platform enables the reproducible study of iPSC-derived neurons and neurogenesis at the single cell level using a miniaturized assay format.

## Materials and Methods

### iPS neuronal progenitor cells (NPCs) and culturing procedures

iPS-derived NPC lines (*i.e.*, 7545-5b and 7753) were derived by and obtained from the University of Virginia Stem Cell Core (Charlottesville, VA) and were previously described^31^. The original fibroblast lines, GM07753 and GM07545 were purchased from the NIGMS Human Genetic Cell Repository at the Coriell Institute for Medical Research (Camden, NJ) by the University of Virginia Stem Cell Core (Charlottesville, VA). The identity of each NPC line was confirmed by immunofluorescence (IF) for nestin and sox2 with minimal to low expression for doublecortin and MAP2. All NPC lines were expanded on Matrigel™-coated dishes in NPC medium [*i.e.*, DMEM-F12 supplemented Glutamax (Gibco, Gaithersburg, MD), 2% B27 without vitamin A (Gibco) and 20 ng/mL thermostable recombinant human Fibroblast Growth Factor (Gibco)] unless otherwise noted. NPC lines were split approximately 1:4 every week (up to passage 15) using Accutase (MilliporeSigma, St. Louis, MO).

### Poly-L-ornithine/laminin coating of tissue culture vessels

All tissue culture ware was coated with poly-L-ornithine (Sigma Aldrich) for 24 hours at 4°C then rinsed with sterile water x 5. Mouse laminin (Gibco) was then added and incubated at 37°C for 4 hours. Poly-L-ornithine/laminin concentrations varied and are as described in the text.

### iPS neuronal differentiation and maturation into neurons under feeder layer-free conditions

NPCs were seeded onto poly-L-ornithine/laminin-coated 6-well and cyclic olefin co-polymer-based 96-well Cell Carrier Ultra microtiter plates (Perkin Elmer, Waltham, MA) at 1×10^5^ and 2×10^3^ cells/well, respectively, unless otherwise noted. For immunofluorescence studies using a Zeiss LSM 700, NPCs were seeded at 1×10^4^ cells per dish onto poly-L-ornithine/laminin coated cyclic olefin co-polymer 35 mm petri dishes (ibidi, Munich, Germany). At 24 h post-seeding, the NPC medium was aspirated and replaced with neuron differentiation medium [DMEM/F12+Glutamax, 2% B27 with vitamin A, 20 ng/mL brain derived neurotrophic factor (Shenandoah Biotechnology, Warwick, PA), 20 ng/mL glial cell derived neurotrophic factor (Shenandoah Biotechnology), 1 mM dibutyryl cyclic adenosine monophosphate (MilliporeSigma), 200 nM ascorbic acid (Millipore Sigma, Burlington, MA), 1 μg/mL mouse laminin (Invitrogen, Carlsbad) and penicillin (100 units/mL)-streptomycin (100 *μ*g/mL) (Thermo Fisher Scientific, Waltham, MA)] to promote cortical neuronal differentiation and maturation unless otherwise noted in the text. BrainPhys^32^ base medium was obtained from StemCell Technologies, (Cambridge, MA) and recombinant human insulin was obtained from Gibco.

### Quantitative polymerase chain reaction (qPCR) assessments using RT2-PCR microarrays

RNA was isolated using an RNAeasy™ kit (Qiagen, Germany) per manufacturer’s instructions. Briefly, iPSC-derived neuron cultures were gently rinsed two times with 1X PBS then were lysed with Buffer RLT supplemented with 1% β-mercaptoethanol. Total RNA was then isolated using an RNAeasy™ column and quantified using a NanoDrop 2000 spectrophotometer (Thermo Fisher Scientific). Total RNA was used to generate cDNA using a RT^2^ first strand synthesis kit (Qiagen). A control cDNA was made using total RNA from normal human adult frontal lobe tissue (BioChain, Newark, CA). RNA concentration was kept consistent within experiments (*i.e.*, 150-500 ng) including controls. SYBR ROX green qPCR reactions were performed in triplicate and data were collected using a BioRad CFX Connect Real-Time PCR Detection System (BioRad, Hercules, CA). Reactions were denatured at 95°C for 10 min then subjected to 95° C for 15 sec and 60° C for 1 min for 40 cycles. Data were analyzed using Microsoft Excel and GraphPad Prism 9.0 (GraphPad, La Jolla, CA). Microarray data were analyzed using Qiagen GeneGlobe. A housekeeping protein control (GAPDH) was used for normalization, using the geometric mean of samples to calculate ΔΔC_T_’s and fold change. The experiment was repeated for a total of three technical replicates. Validated primer sets were obtained from Qiagen.

### Western blotting

Whole cell lysates were generated using M-PER™ (ThermoFisher) with 1X Halt™ protease and phosphatase inhibitor (ThermoFisher), and were electrophoresed using 4-12% SDS-PAGE gels (BioRad). Proteins were transferred, per manufacturer’s instructions, to PVDF membranes using a BioRad Trans-Blot Turbo Transfer System. The membrane was allowed to dry at RT for one hour, then was activated with methanol for 2 min and rinsed briefly with water. Membranes were then incubated in Intercept blocking buffer (LI-COR, Lincoln, NE) overnight with rocking at 4°C. The membranes were then incubated overnight at 4° C in a primary antibody diluted to 1 μg/mL in Intercept blocking buffer. Blots were washed three times for five minutes with 1XTBS, then incubated for 2 hours at 4° C in either DyLight 800 goat anti-rabbit IgG (Invitrogen) or DyLight 800 rabbit anti-mouse IgG (Rockland Immunochemicals, Limerick, PA) secondary antibodies diluted to 100 ng/mL in Intercept blocking buffer. Membranes were washed three times for five minutes with TBS, then imaged on a LI-COR Odyssey CLx Near-Infrared Imaging System. Band signals were quantified using Image Studio software (LI-COR). Lane normalization factors for each sample were determined per LI-COR recommendations by dividing the GAPDH signal for a sample by the highest value signal of GAPDH on the blot. The target protein signal was then normalized to GAPDH by dividing the raw signal value by the lane normalization factor.

### Amyloid-β_1-42_ ELISA

An amyloid-*β*_1-42_ ELISA kit was obtained from R&D Systems (Minneapolis, MN) and performed using manufacturer’s instructions. For cell supernatant harvesting, maintenance medium was aspirated and replaced with 2 mL of new medium. Cultures were incubated for 24 h and supernatants were harvested. After clarification by centrifugation at 4° C, cell supernatants were stored at −80° C until evaluation.

### Electrophysiological assessments of iPSC-derived neurons

Electrophysiological assessments were performed as described^33^. Briefly, 7753 NPCs (1×10^5^) were seeded onto cyclic olefin co-polymer coverslips (ibidi GmbH, Munich, Germany) that were cut into 2.5 cm diameter circles and coated with 50 μg/mL poly-L-ornithine and 25 μg/mL laminin. Cells were allowed to differentiate for 4 weeks under 5% O_2_. All electrophysiology recordings were performed and analyzed using pClamp™ and Clampfit™ software (Molecular Devices, LLC, San Jose, CA, USA). During the recording procedure, cultures were perfused with BrainPhys™ medium. Action potentials were manually measured from the trough of each spike to the peak and events greater than 10 mV in amplitude were considered action potentials.

### Cell fixation, IF and image acquisition

Cells were fixed using 1% paraformaldehyde (PFA, Thermo Scientific) in 1X Dulbecco’s phosphate buffered saline (DPBS) with calcium and magnesium and supplemented with 100 nM DAPI (Life Technologies, Eugene, OR). After 3 minutes at room temperature, PFA was replaced with ice-cold 100% methanol (Fisher) and incubated for 3 min at room temperature. Four times the volume of DPBS was then added to methanol containing wells, immediately removed and then replaced with DPBS. This DPBS replacement step was repeated and the plate was sealed with aluminum film (Axygen Biosciences, Corning, NY) for storage at 4° C until immunofluorescence (IF) staining.

For IF antibody staining, DPBS was removed and cells were incubated with 0.3% Triton-X in 1X DPBS for 10 minutes, and then were gently washed twice with 1X DPBS for 5 minutes each. After the final wash, DPBS was removed and replaced with 300 mM glycine in 1X DPBS for 20 minutes. After quenching, the solution was removed and replaced with 1% Fish Skin Gelatin (Millipore Sigma) blocking buffer in DPBS containing primary antibody. Primary antibody staining was carried out at 4°C in a humidified chamber for 16-18 hours. Antibody solution was then removed and wells were washed 3 times with 1X DPBS for 5 minutes each. If primary antibodies were not directly conjugated with fluorophore, conjugated secondary antibodies in 1% Fish Skin Gelatin blocking buffer were added and incubated for 24 hours at 4°C, protected from light in a humidified chamber. Conjugated secondary antibody solution was then removed, wells were washed three times for 5 minutes each at room temperature, and 1X DPBS storage buffer was added for imaging.

The Perkin Elmer Operetta high content imaging system, located in the University of Virginia Advanced Microscopy Core Facility, was used for image acquisition unless otherwise noted. Perkin-Elmer Harmony 4.1 software was used to analyze acquired images.

### Antibodies

Antibodies were obtained and used as described in Supplemental Table 1.

### Statistical analysis

Data were analyzed using GraphPad Prism 9.0. Data are presented as mean ± standard deviation or standard error of the mean. *p* Values were calculated with Student’s *t*-test for comparisons involving 2 groups or one-way or two-way analysis of variance for comparisons involving >2 groups. *p* < 0.05 was considered statistically significant.

## Results

### Tracking iPSC-derived neuron maturation at environmental O_2_ (21%)

We differentiated NPCs derived from healthy individuals under feeder layer-free conditions in environmental O_2_ (21%) for 12 weeks. At 2 week intervals, we isolated total RNA for qPCR and protein for western blotting. Calretinin, the first post-mitotic protein expressed in “early” mature neurons^34^, was expressed ~6-8 weeks post-differentiation of 7753 cells (Figure 1, panels A, D). Detection of βIII-tubulin at just 2 weeks post-differentiation confirmed the presence of cells on the neuron differentiation pathway (Figure 1, panel A; Supplemental Figure 1). Similar results were observed using another NPC line, 7545-5b, derived from a healthy patient (Supplemental Figure 2). In contrast, gene expression of other neuronal maturity markers, including Calbindin, NeuN, MAP2 and Tau, rose appreciably from initially low levels only after 4-10 weeks of differentiation (Figure 1, panels A and D). Moreover, while the NPC marker, nestin, and the cycling cell markers, PCNA and cyclin D1, decreased with time (Figure 1, panels A, B, and C), their persistent detection at 12 weeks post-differentiation imply that subpopulations within the iPSC-derived neuronal cultures were actively proliferating and differentiating for at least 12 weeks.

**Figure 1.**
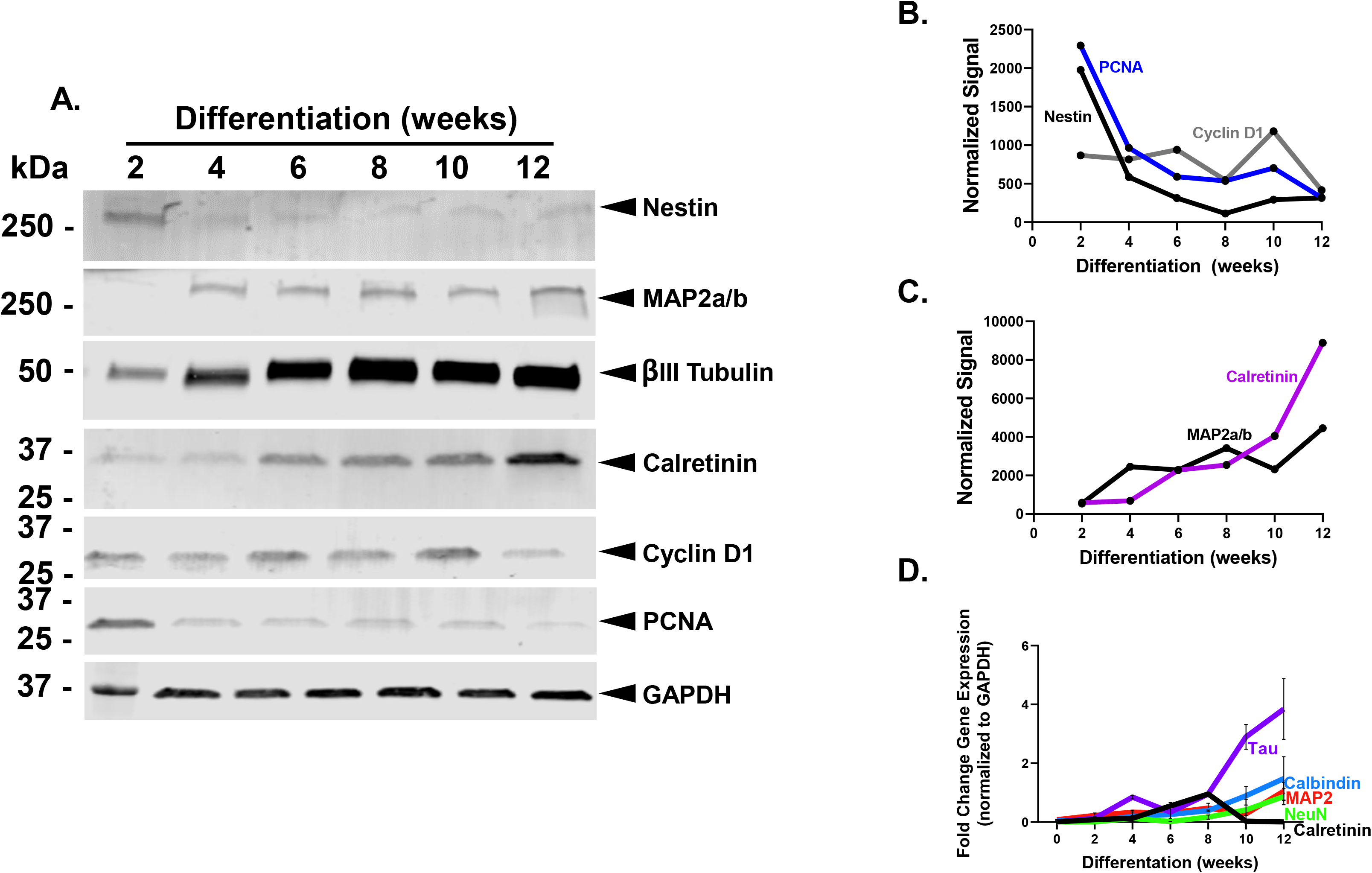
Expression of neuronal differentiation and maturation markers under environmental (21%) O_2_. Neuronal progenitor cells (*i.e.*, 7753) from healthy patients were differentiated under 21% O_2_ using feeder layer-free conditions. RNA and protein lysates were isolated every two weeks for 12 weeks. Panel A, western blot assessments for neuronal maturation markers and cell cycle associated signaling effectors. Panels B and C, GAPDH normalized signals for select proteins. Panel D, qPCR time course data for neuronal maturation markers. N=3, mean ± SD.

### Enhancement of iPSC-derived neuron maturation under physiological O_2_ (5%)

Since normal oxygen levels in the human brain are ~5%^35^, environmental O_2_ culturing conditions (~21%) may not accurately reflect the brain physiological milieu. To determine the effects of physiological oxygen tension (5%) on the differentiation and maturation of iPSC-derived feeder layer-free neuronal cultures, we tracked calretinin gene expression over an 8-week period to resolve the “calretinin spike.” Thus, normal, healthy NPCs were differentiated and matured as described^36^ under environmental (21%) or physiological (5%) O_2_ tension using the same ECM composition (5 μg/mL laminin and 10 μg/mL poly-L-ornithine [PLO]), medium formulation and medium change schedule. RNA was collected at the specified times and resulting qPCR data indicated that the calretinin spike occurred at 6 weeks post-differentiation under both O_2_ culturing conditions, but the magnitude of the calretinin spike was significantly more pronounced with 5% O_2_ (Figure 2, panel A). Calretinin protein expression was confirmed by western blot (Figure 2, panels B and C). As expected, nestin protein expression declined with neuronal maturation (Figure 2, panels B and D), but the decrease was more pronounced under 5% O_2_. The lower O_2_ level also was associated with less Aβ_42_ peptide accumulation in the medium (Supplemental Figure 3), which may enhance neuron viability in long term culture^37^. Overall, these data favor the hypothesis that iPSC-derived neurons mature more quickly in 5% versus 21% O_2_. Moreover, they underscore the ability of physicochemical cues to modulate progenitor cell fate and function^38^, which may be particularly critical for NPC-derived model systems.

**Figure 2.**
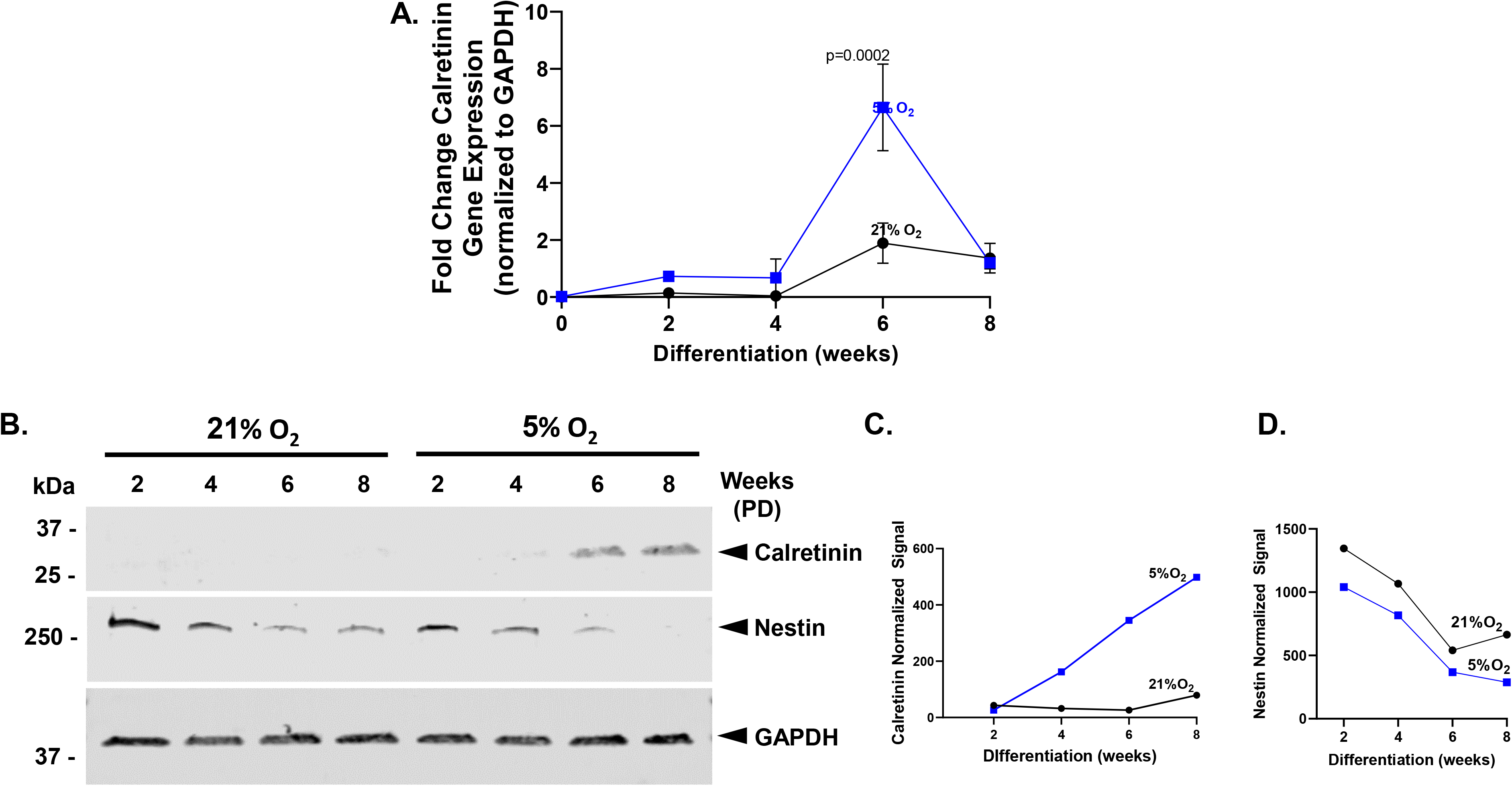
Enhanced calretinin gene and protein expression under 5% O_2_ and feeder layer-free culturing conditions. Neuronal progenitor cells (line 7753) derived from apparently healthy patients were matured under environmental (●, 21%) vs. physiological (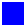, 5%) O_2_ in 6 well plates. Panel A, Calretinin gene expression was quantified by qPCR. Statistics were performed using two-way ANOVA. N=3. Panel B, Calretinin, nestin and GAPDH protein expression as detected by western blot. PD=post-differentiation. Panels C and D, Quantification of calretinin and nestin western blots. Proteins were normalized using GAPDH.

### Shifting the “calretinin spike” through modifications of extracellular cues

The six-week time course for the switch to a more mature neuronal phenotype is consistent with previous studies^39^; however, the extended maturation duration required for derivation of mature neuron populations is experimentally limiting, especially for miniaturized assay formats. To further accelerate iPSC neuron differentiation and maturation, we modified the composition of the ECM and culture media. Using 50 μg/mL poly-L-ornithine (PLO)/15 μg/mL laminin as the ECM and a gradual transition to a 100% BrainPhys^32^-based differentiation medium (2% B27 with insulin, 20 ng/mL BDNF, 20 ng/mL GDNF, 0.2 μM ascorbic acid, 1 μg/mL laminin, 1 mM cAMP, and Pen (5,000 units/mL)-Strep (5,000 μg/mL)) over 48 h, the “calretinin spike” shifted from 6 weeks (Figure 2, panel A) to 4 weeks (Figure 3, panel A). Moreover, increasing the laminin concentration to 25 μg/mL further shifted the “calretinin spike” to two weeks (Figure 3, panel A) indicating an acceleration of the differentiation/maturation of iPSC-derived neurons through manipulation of the extracellular environment. The magnitude of the “calretinin spike” differs between the two ECM conditions, potentially reflecting the existence of specific neuronal progenitor subpopulations which may be more receptive to the extracellular cues leading to differentiation/maturation. Nonetheless, western blotting and immunofluorescence studies confirmed the expression of Tau and NeuN at four weeks post-differentiation (Figure 3, panels B and C) verifying that feeder layer-free culturing conditions can support differentiation of mature iPSC-derived neurons.

**Figure 3.**
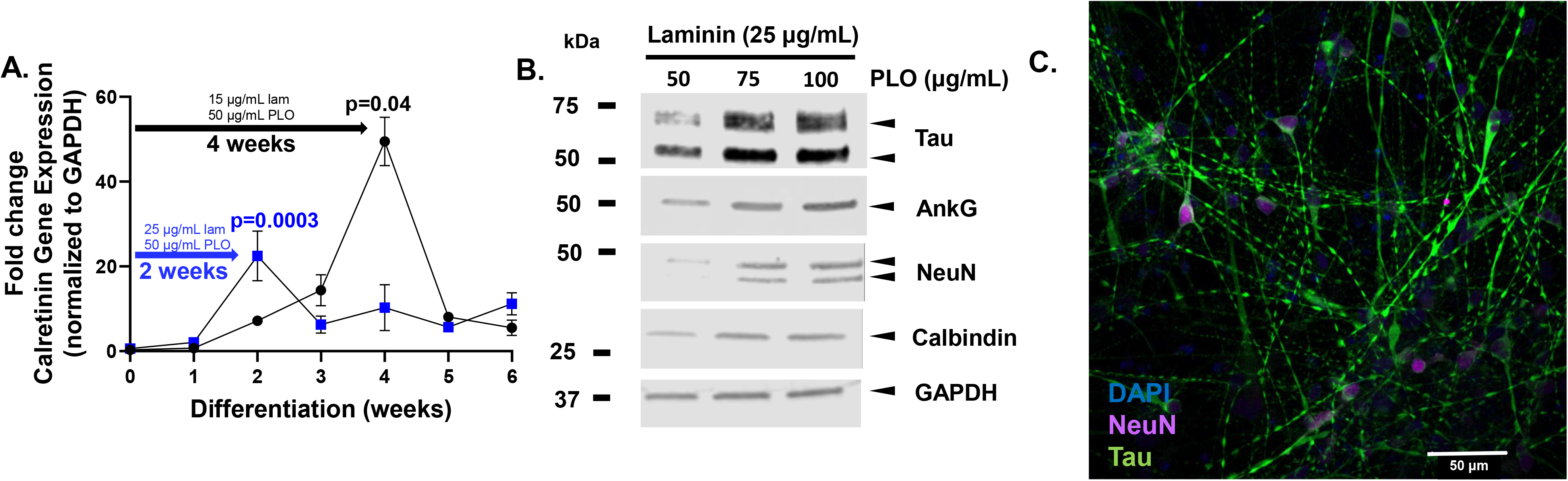
Extracellular cues shift the “calretinin spike” during feeder layer free iPSC-neuronal differentiation and maturation. Panel A, iPSC-derived neurons from apparently healthy patients were matured under physiological (5%) O_2_ in 6 well plates coated with (●) 15 ug/mL laminin and 50 ug/mL PLO or 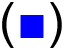 25 ug/mL laminin and 50 ug/mL PLO. Calretinin gene expression was quantified by qPCR. Statistics were performed using two-way ANOVA. N=3. Panel B, Expression of proteins associated with neuronal maturation. Panel C, Expression of Tau and NeuN in four-week feeder layer free iPSC-derived neuron cultures as detected by immunofluorescence.

### Identification and quantification of the “neurogenesis burst”

Using the optimized ECM and culture medium transitions, we miniaturized the assay format to a 96 well plate to permit high content imaging (HCI). We initially seeded 1,500 NPCs per well of a 96 well plate and allowed them to differentiate and mature over a 6-week time course with medium changes occurring twice weekly. Each week a microtiter plate was fixed and stained with the nuclear dye, 4′,6-diamidino-2-phenylindole (DAPI), and total DAPI+ cells (*i.e.*, nuclei) were quantified. This showed that between weeks 2 and 3 in differentiation medium, the NPC cultures undergo a “neurogenesis burst” characterized by a rapid increase in total DAPI+ cells (Figure 4, panels A and B). By week 4 post-differentiation, each well contained ~15,000 neurons, representing a 10-fold increase in cell number versus NPC seeding. After 3-weeks post-differentiation, DAPI+ cell number plateaued suggesting a cessation in the proliferation of early/intermediate neuronal progenitor cells. By 4 weeks, the expression of markers of a more mature neuronal phenotype (*i.e.*, Calbindin+, NeuN+) was detectable by immunofluorescence in a subpopulation of DAPI+ cells (Figure 5, panel A-C). Specifically, data showed that at 4-weeks post-differentiation 30-40% of DAPI+ cells stained positive for Calbindin and between 10-15% of Calbindin+ cells also expressed NeuN (Figure 5, panels A and B). Although these percentages are low for “advanced maturation”, we note that the maturation of the cell cultures is asynchronous and with additional incubation, the Calbindin+/NeuN+ subpopulation would be expected to increase. Moreover, it is likely not all of the cells will progress to mature neurons (*i.e.*, as defined by Calbindin+/NeuN+) under feeder layer-free culturing conditions.

**Figure 4.**
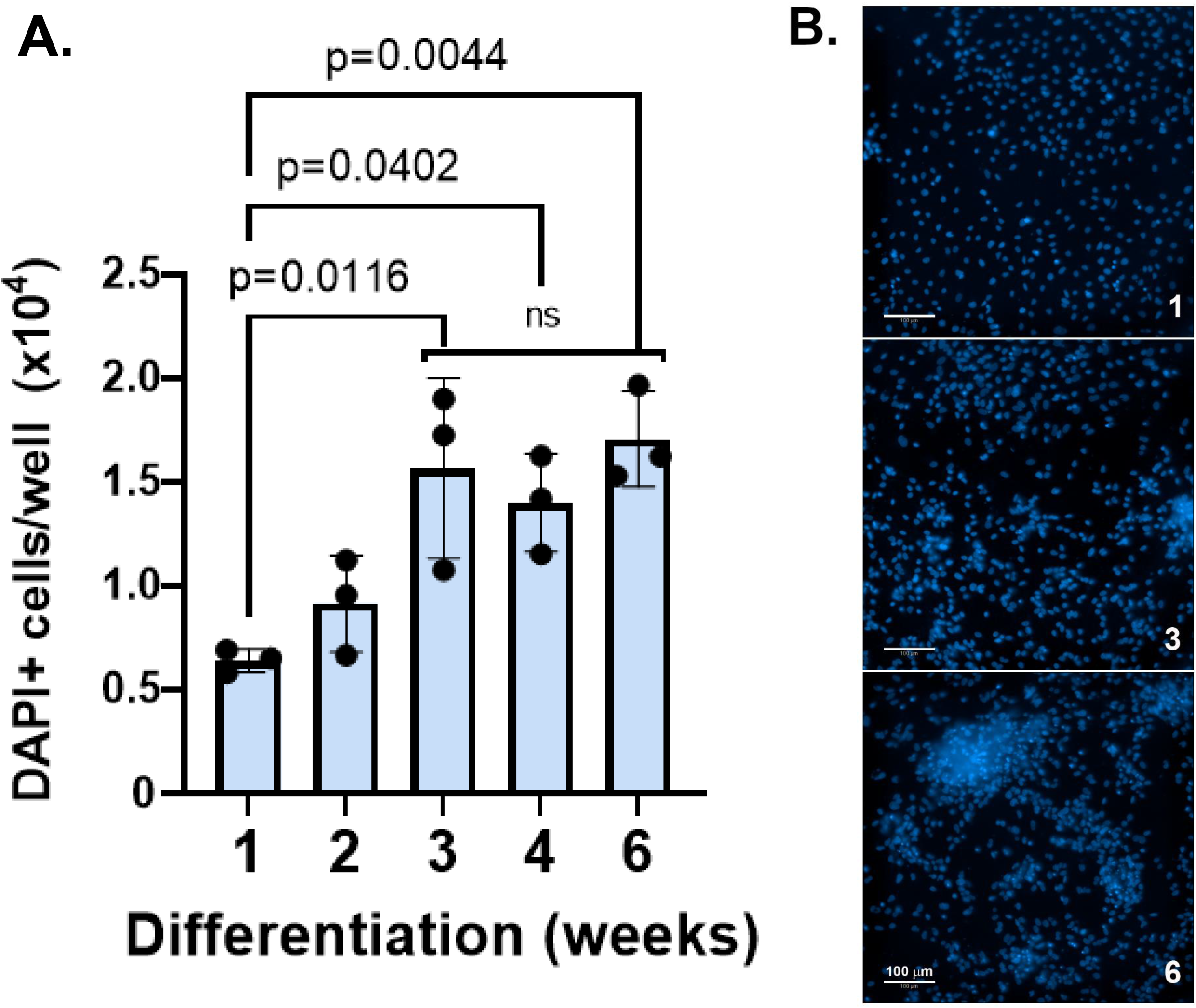
Quantification of the “neurogenesis burst” over a six-week time course under feeder layer free conditions. 1-6 week iPSC-derived neuron cultures were stained with DAPI to track the “neurogenesis burst.” Panel A, DAPI+ nuclei were quantified in 25 fields of view per well of a 96 well plate comprising differentiating neuronal populations. NPCs were seeded at 1,500 cells per well on 25 μg/mL laminin and 50 μg/mL poly-L-lysine. N=3 technical replicates. One-way ANOVA. ns, not significant. Panel B, DAPI+ images of fields of view from 1, 3 and 6 week differentiating neuronal cultures.

**Figure 5.**
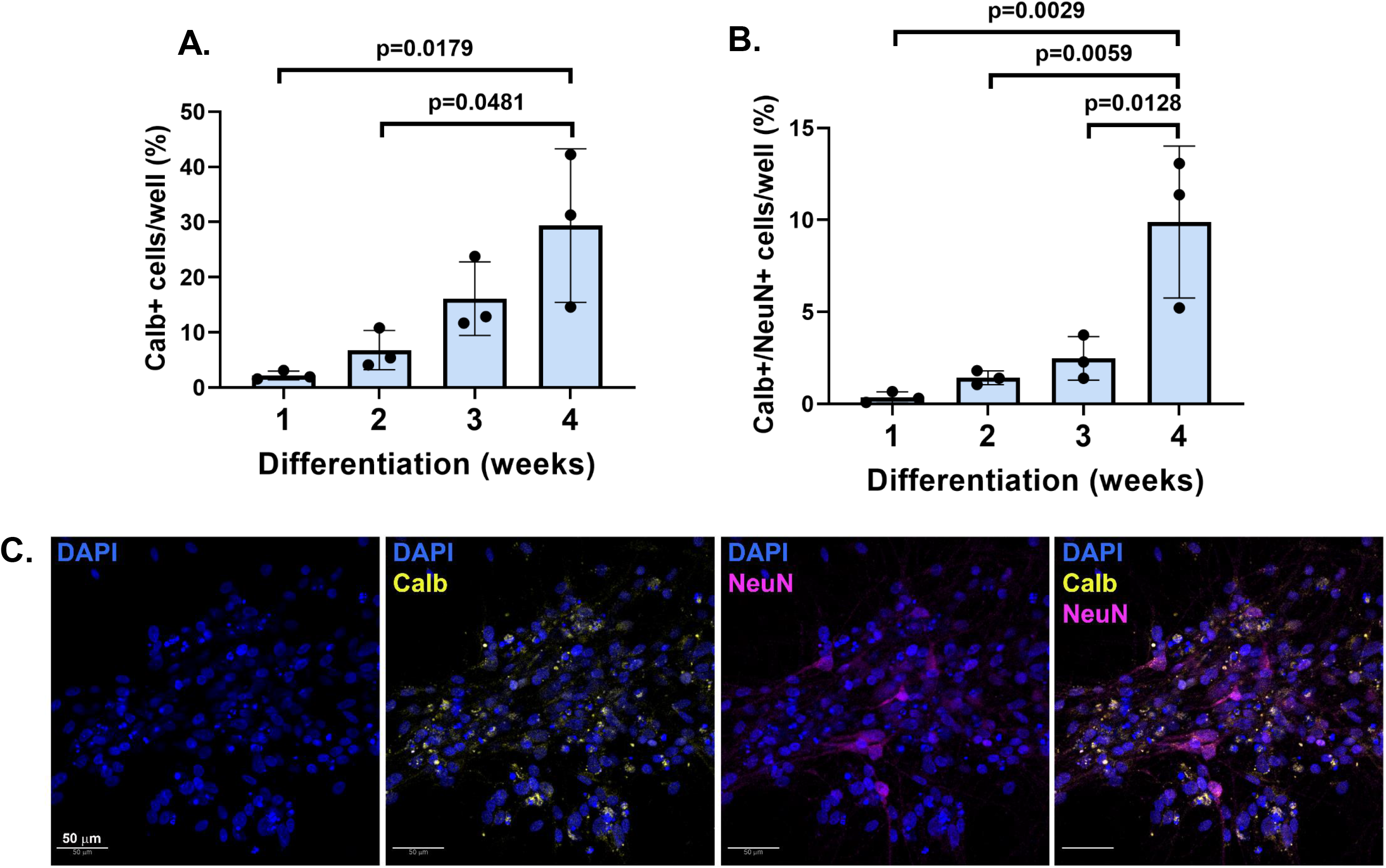
Immunofluorescence detection of Calbindin and NeuN expression by four-week iPSC-derived neuronal cultures. 1-4 week iPSC-derived neuron cultures were stained with DAPI, Calbindin and NeuN to identify mature neurons. DAPI+ nuclei were identified and quantified using 25 fields per well. Panel A, Calbindin+ cells were identified using anti-Calbindin-Alexa-488 (Cell Signaling, #88831). Panel B, NeuN+ cells were identified using anti-NeuN-Alexa-fluor-647 (Millipore-Sigma, HPA030790). Colocalized signals were quantified to determine the percent Calbindin+/NeuN+ cells. N=3 technical replicates. One-way ANOVA. Panel C, Immunofluorescence images of Calbindin+, NeuN+ and Calbindin+/NeuN+ cells.

### Cessation of the “neurogenesis burst” coincides with the acquisition of a non-cycling phenotype

The neuronal population evaluated at 4-weeks post-differentiation was also non-cycling, supporting the acquisition of a more mature neuronal phenotype. Specifically, on a population level, qPCR microarray data showed that the expression of cyclin D1 and Ki67, two biomarkers for proliferation, was significantly downregulated (Figure 6, panels A and B). Cyclin D1 also plays a key role in regulating iPSC-derived neuronal differentiation^40^. Downregulation of cyclin D1 and Ki67 gene expression in these asynchronously differentiating cultures coincided with upregulated expression of genes associated with neuronal maturation including CDK7 (*e.g.*, synaptic plasticity^41^), CDK5RAP1 (*i.e.*, neuronal differentiation^42^), and CASP3 (*e.g.*, synaptic activity, dendrite pruning^43^). Moreover, we detected the expression of genes associated with normal post-mitotic neurons (*e.g.*, CCNH^44^) or those that are involved with neuronal cell maintenance (*i.e.*, ATR^45^). Although most commonly associated with senescence^46^, CDKN2B inhibits iPSC division and regulates tissue remodeling^46–47^. Nonetheless, Ki67 gene expression data were confirmed on a single cell basis using immunofluorescence (<7% expression, Figure 6, panels C and D) in our miniaturized format with data showing that by 3-4-weeks post-differentiation Ki67 expression levels were significantly decreased (Figure 6, panels C and D, Supplemental Figure 4). Within the Calbindin+/NeuN+ subpopulation of cells, Ki67 positivity was <~2% (data not shown).

**Figure 6.**
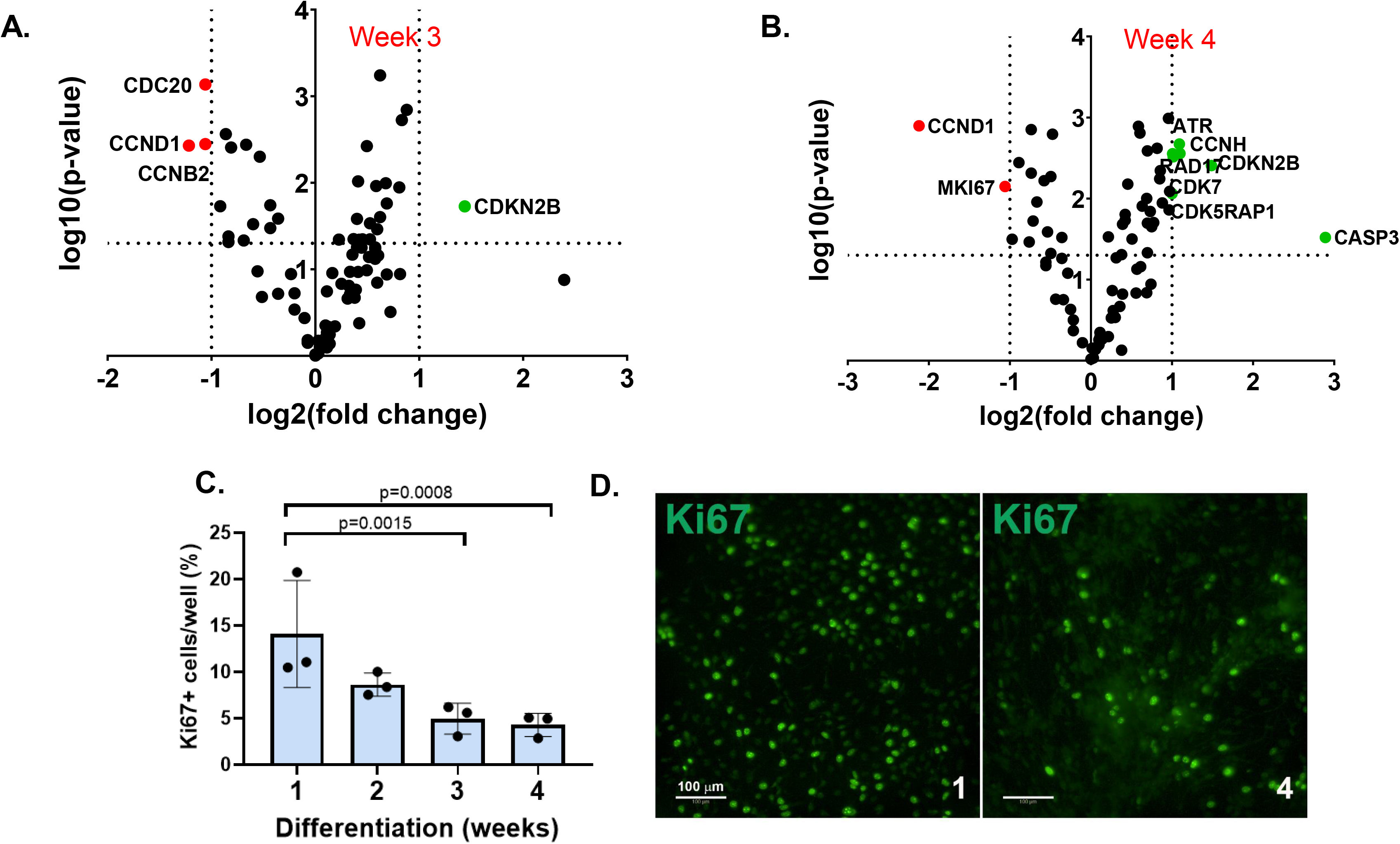
Decrease in Cyclin D1 and Ki67 gene expression with iPSC-derived neuronal differentiation and maturation under feeder layer-free conditions. Panels A and B, qPCR microarray data comparing cell cycle associated gene expression in week 3 (A) and 4 (B) iPSC-derived neuronal cultures. Panel C, Ki67+ positive cells were quantified over a 4 week culturing period. Twenty-five fields of view were captured per well (N=3) and Ki67 was detected using anti-Ki67-Alexa-488 conjugate. One-way ANOVA. Panel D, representative field of view of Ki67 positivity in week 1 versus week 4 iPSC-derived neuronal cultures.

Electrophysiological assessments of our 4-week cultures also demonstrated that feeder layer-free iPSC-derived neurons were electrically excitable. Specifically, recorded neurons maintained the relatively depolarized resting membrane potential of ~-40mV and generated action potentials in response to current injection (Figure 7). The amplitude of the action potential was generally smaller during depolarizing current injection than during hyperpolarizing current injection, suggesting that voltage hyperpolarization-dependent sodium channel de-inactivation was necessary for the generation of robust, large amplitude action potentials. This observation, in combination with the depolarized resting voltage, indicates that while 4-week old neurons are generally mature and electrically-excitable, these neurons nonetheless lack a pronounced resting potassium conductance found in adult neurons.

**Figure 7.**
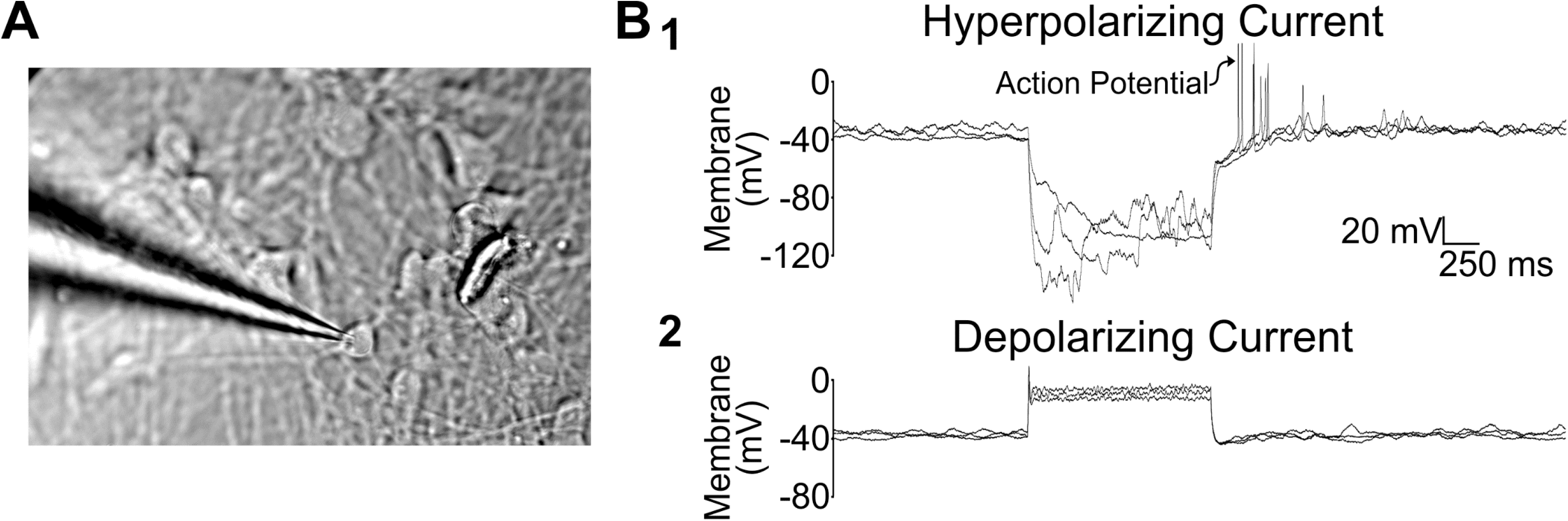
iPSC-derived neurons cultured under feeder layer-free conditions conduct action potentials. Panel A, Patch clamp recording of a 4 week iPSC-derived neuron. Panel B, Electrophysiological assessments of iPSC-derived neurons at 4 weeks post-differentiation indicate they conduct action potentials. Panel B, The neuron’s resting membrane potential is around −40 mV (versus ~60 mV for most adult neurons). In response to hyperpolarizing current injection (B1), the neuron produced multiple action potentials characterized by a large amplitude. These action potentials were typically +60 mV in amplitude and persisted for a duration of ~4-5 msec. In response to positive current (B2), the recorded neuron produced a single action potential of moderate amplitude. The discrepancy in action potential amplitude observed during hyperpolarizing versus depolarizing current injection is consistent with the hypothesis that the relatively depolarized resting membrane potential of recorded neurons promotes steady-state sodium channel inactivation, which can be removed by hyperpolarizing the voltage of the neuron.

## Discussion

Dysfunctional neurogenesis is evident in multiple neurological *(e.g.*, epilepsy, schizophrenia, depression, anxiety, addiction)^48–50^ and neurodegenerative disorders (*e.g.*, Parkinson’s Disease, Alzheimer’s Disease, Huntington’s Disease, Prion Disease)^51–52^. Unfortunately, deficits in neurogenesis, which lead to neuronal decline, most likely initiate many years prior to the manifestation of cognitive, memory and/or behavioral issues underscoring their enduring repercussions. Hence, there is intense interest to generate new model systems to study and promote neurogenesis.

iPSC-derived neurons offer physiologically relevant cell populations to study neurological processes, including neurogenesis. In contrast, “iNeurons”, which are directly converted from adult fibroblasts and retain adult, rather than embryonic epigenetic signatures, bypass the proliferating multi-functional progenitor (*i.e.*, NPC) stage and, hence, are more limiting as models of early events in neurogenesis^53^. Regardless, the prolonged differentiation and maturation durations present experimental challenges and limit the usefulness of iPSC-derived cell populations (as well as iNeurons) especially in miniaturized formats. Through minimal modification of extracellular cues (underlying ECM, medium composition^32^/transitions and culturing conditions [5% O_2_]), however, we determined that iPSC-derived differentiation and maturation of a feeder layer-free system can be accelerated suggesting that with the incorporation of more bio-complex and -relevant extracellular signals or cues, neurogenesis may be further advanced and regulated without the need for an astrocyte feeder layer.

Our current ECM has two primary components, laminin and poly-D/L-ornithine (PLO), both of which frequently are used in ECMs for iPSC-derived neuronal differentiation and maturation^54–55^. Although laminin is physiologically relevant, it represents a single ECM component whereas the brain ECM consists of multiple macromolecules^56^, including glycosaminoglycans, proteoglycans, glycoproteins and fibrous proteins. Moreover, the composition of the brain ECM changes during neurodevelopment in order to provide the extracellular/environmental cues necessary to control neurogenesis^57^ suggesting that the brain ECM is more fluid than previously surmised. Unsurprisingly, recent studies have shown that neural development is accelerated when using ECM from decellularized brain tissue^57^, which comprises ECM proteins from all cell populations within the brain. However, as is the case for Matrigel, decellularized brain ECM it is subject to substantial batch-to-batch variation. Moreover, studies examining the role of a more bio-complex ECM in neuronal development tend to be in lower density formats^58^ (6-24 well plates) while also relying on astrocyte feeder layers to derive mature neurons, two conditions that are not conducive to single cell analyses.

Normal tissue O_2_ levels are variable but generally fall in a range of 3–9%^59^, which is less than atmospheric O_2_ (*i.e.*, 21%) in the air we breathe and what is commonly used in cell culturing activities. Studies have also shown that physiological O_2_ levels are not only critical for neuronal stem cell self-renewal, but also limit spontaneous differentiation^60^. Unfortunately, the use of physiological O_2_ for differentiating and maintaining iPSC-derived neurons is only sporadically mentioned in the literature^58^. Moreover, the prevalence of atmospheric O_2_ usage in neuronal studies may contribute to the lack of reproducibility, at least with respect to iPSC-derived neuronal differentiation and maturation.

Additional modifications to our iPSC-derived neuron culturing medium and transitions limited the asynchronous proliferation of early progenitor cells (Supplemental Figure 5). As a result, we were able to miniaturize the iPSC-derived neuronal differentiation and maturation process. Through the use of a HCI platform, we identified and quantified the “neurogenesis burst” associated with the rapid proliferation of progenitor cells, which leads to the development of mature iPSC-derived neurons as identified by Calbindin and NeuN co-expression.

**Table 1.** Antibodies used for immunofluorescence and western blotting.

| **Protein target** | **Vendor** | **Catalog number** | **Concentration** | **Application** |
| --- | --- | --- | --- | --- |
|  |  |  | (µg/mL) |  |
| **Calbindin** | Novus | NBP2-67200 | 1 µg/mL | WB |
| **Calretinin** | Cell Signaling | 17114S | 1 µg/mL | WB |
| **CyclinD1** | Cell Signaling | 55506S | 1 µg/mL | WB |
| **Doublecortin** | Cell Signaling | 4604S | 1 µg/mL | WB |
| **GAPDH** | Cell Signaling | 2118S | 1 µg/mL | WB |
| **Ki67-Alexa 488** | Cell Signaling | 11882 | 0.5 µg/mL | IF |
| **MAP2** | Cell Signaling | 4542S | 1 µg/mL | WB |
| **Nestin** | Cell Signaling | 33475S | 1 µg/mL | WB |
| **NeuN** | Cell Signaling | 12943S | 1 µg/mL | WB |
| **NeuN (RBFOX3)** | Sigma | HPA030790 | 0.4 µg/mL | IF |
| **PCNA** | Cell Signaling | 2586S | 1 µg/mL | WB |
| **Tau** | Cell Signaling | 4019S | 1 µg/mL | WB, IF |
| **Tau** | Cell Signaling | 46687 | 4 µg/mL | IF |
| **Tuj1 (βIII tubulin, TUBB3)** | From Bloom lab | N/A | 1 µg/mL | WB |
| **Synaptophysin** | Abcam | ab8049 | 1:1000 dilution | WB, IF |
| **Anti-Mouse Alexa 488** | Cell Signaling | 4408 | 2-4 µg/mL | IF |
| **Anti-Mouse Alexa 647** | Cell Signaling | 4410 | 2-4 µg/mL | IF |
| **Anti-Rabbit Alexa 488** | Invitrogen | A11008 | 2-4 µg/mL | IF |
| **Anti-Rabbit Alexa 647** | Cell Signaling | 4414 | 2-4 µg/mL | IF |

## Acknowledgements

The authors thank Dr. Stacey Criswell (University of Virginia Advanced Microscopy Facility) for her assistance with image capture on the Zeiss LSM 700. This research was made possible through funding from the National Institutes of Health (R01 AG063400, ERS), the Cure Alzheimer’s Fund (ERS, JSL, GSB), the Owens Family Foundation (JSL, GSB), the Rick Sharp Alzheimer’s Disease Foundation (JSL) and the Fiske Drug Discovery Laboratory.

## Notes

### Competing Interest Statement

The authors have declared no competing interest.

