## Supplemental Data for "Extracellular cues accelerate neurogenesis of induced pluripotent stem cell derived neurons"

### Supplemental Figures & Tables

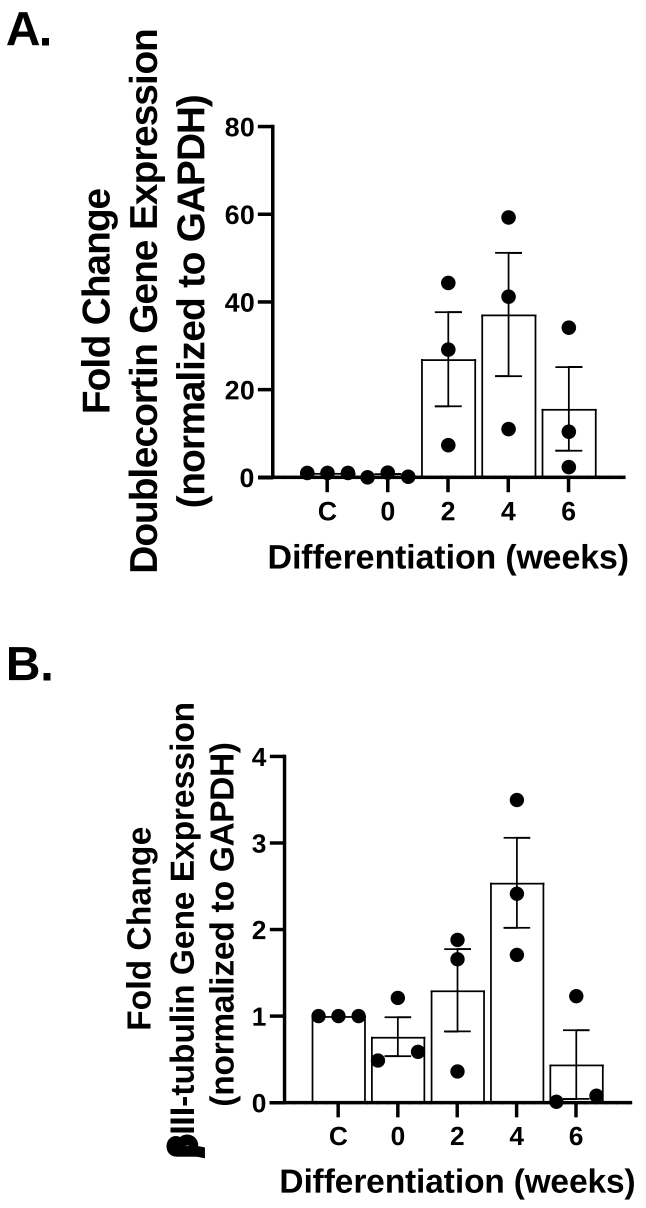


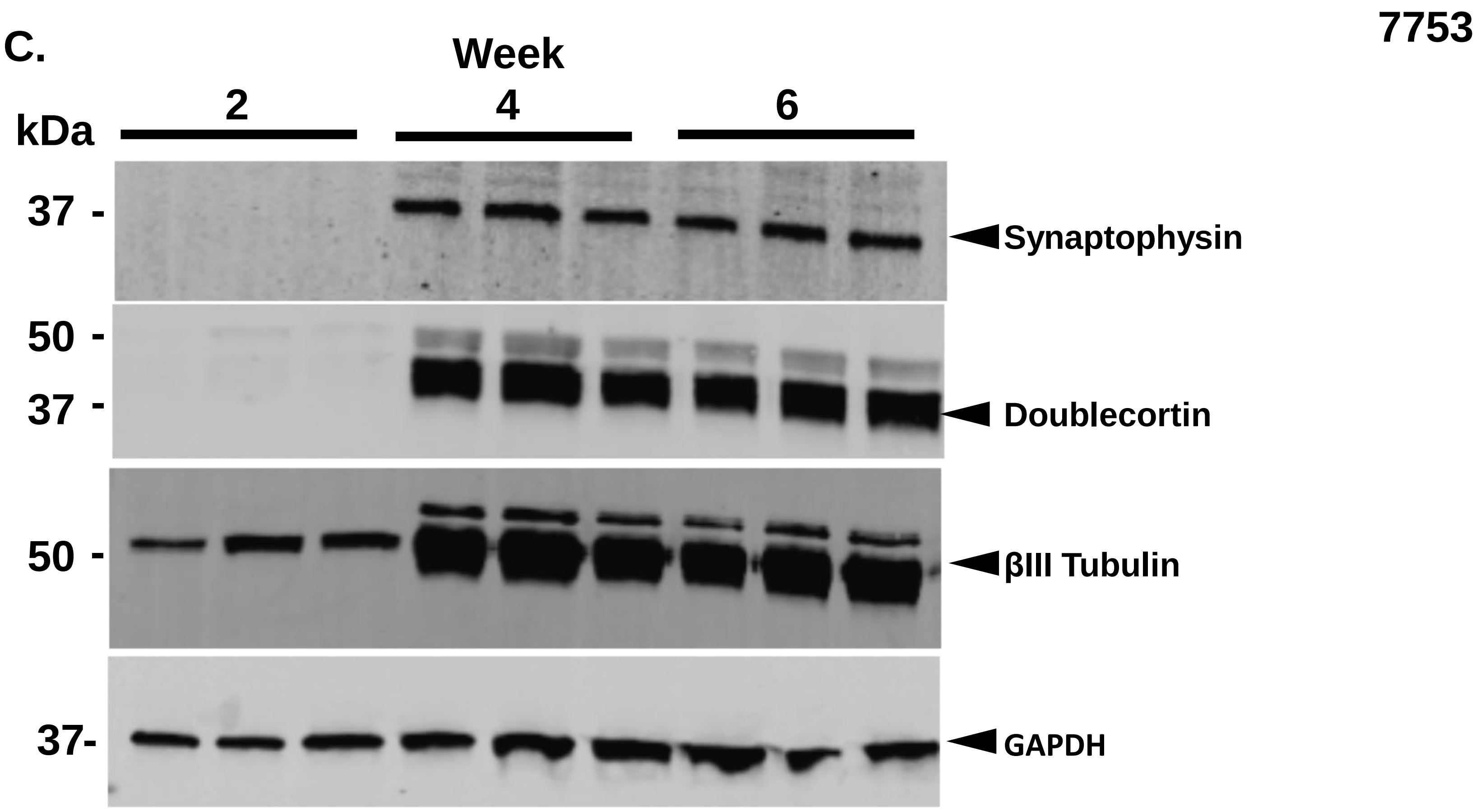


**Expression of neuronal differentiation and maturation markers in the 7753 NPC line under environmental (21%) $O_2$.** Neuronal progenitor cells (*i.e.*, 7753) derived from healthy patients were differentiated at 21% $O_2$ using feeder layer-free conditions. RNA and protein lysates were isolated every 2 weeks for 12 weeks. Panels A and B, doublecortin and βIII-tubulin gene expression as quantified by qPCR. N=3, mean ± SD. Panel C, Western blot assessments for neuronal maturation markers. Week=post-differentiation induction.

#### Supplemental Figure 1

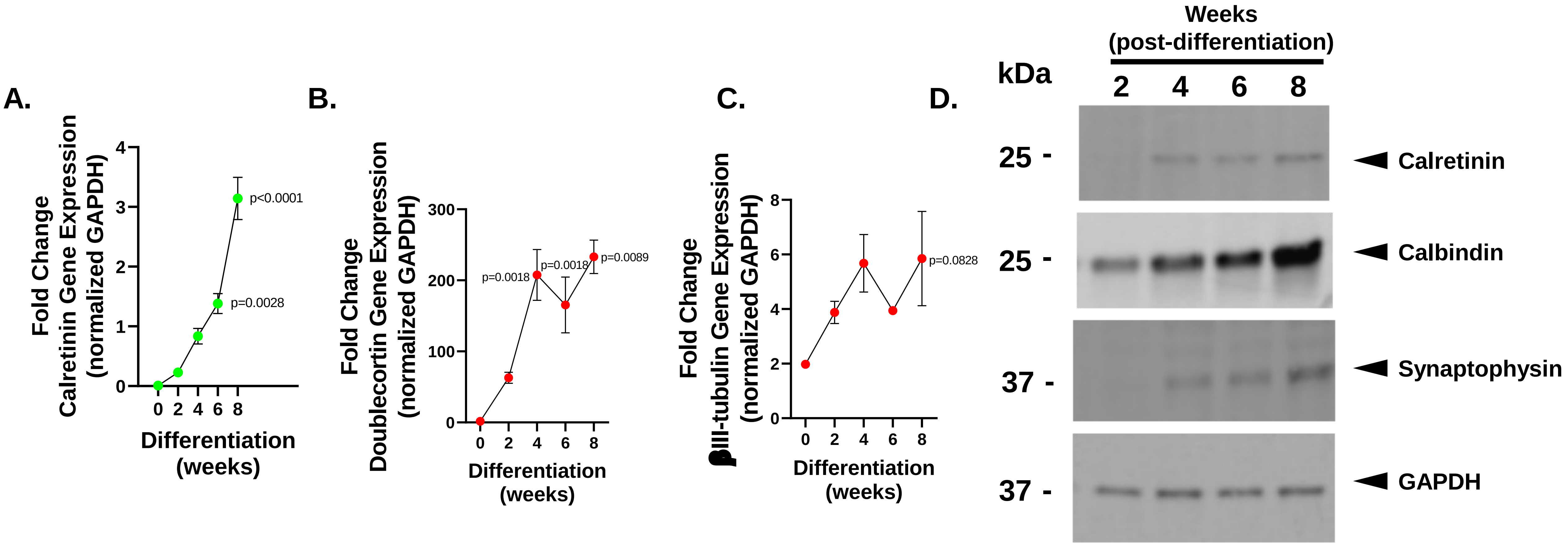


**Expression of neuronal differentiation and maturation markers in the 7545 NPC line under environmental (21%) $O_2$.** Neuronal progenitor cells (line 7545) derived from healthy patients were differentiated under 21% $O_2$ using feeder layer-free conditions. RNA and protein lysates were isolated every 2 weeks for 8 weeks. Differentiating and maturing NPCs displayed a slightly different kinetics of expression of neuronal maturation markers. Panels A-C, qPCR time course data for neuronal maturation markers. N=3, mean ± SD. Panel D, western blot assessments for neuronal maturation markers.

#### Supplemental Figure 2

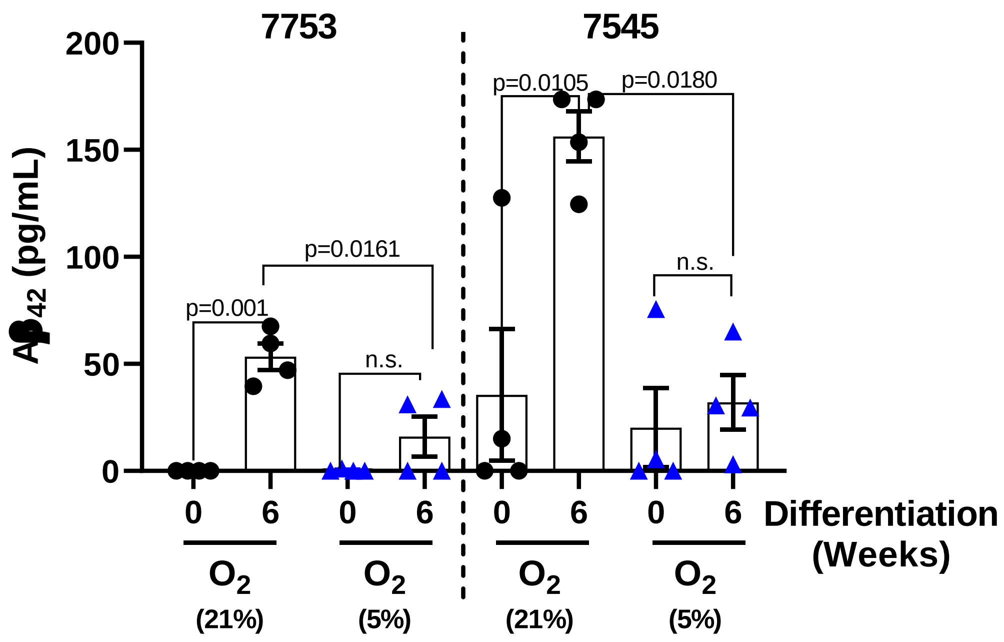


**Physiological $O_2$ levels suppress $A\beta_{42}$ secretion by healthy NPCs during differentiation and maturation.** Normal, healthy patient-derived neuronal progenitor cells (lines 7753, 7545) were seeded into six-well plates coated with 5 µg/mL laminin and 10 µg/mL poly-L-lysine and differentiated/matured as described in text using physiological $O_2$ (5%) versus environmental $O_2$ (21%). Medium was changed at the designated timepoints and 24 hours later, culture supernatants were isolated and an $A\beta_{42}$ ELISA was performed according to manufacturer's instructions. N=4, mean± SD. Data analyzed using one-way ANOVA. ●, Environmental $O_2$ (21%). ▲, Physiological $O_2$ (5%).

### Supplemental Figure 3

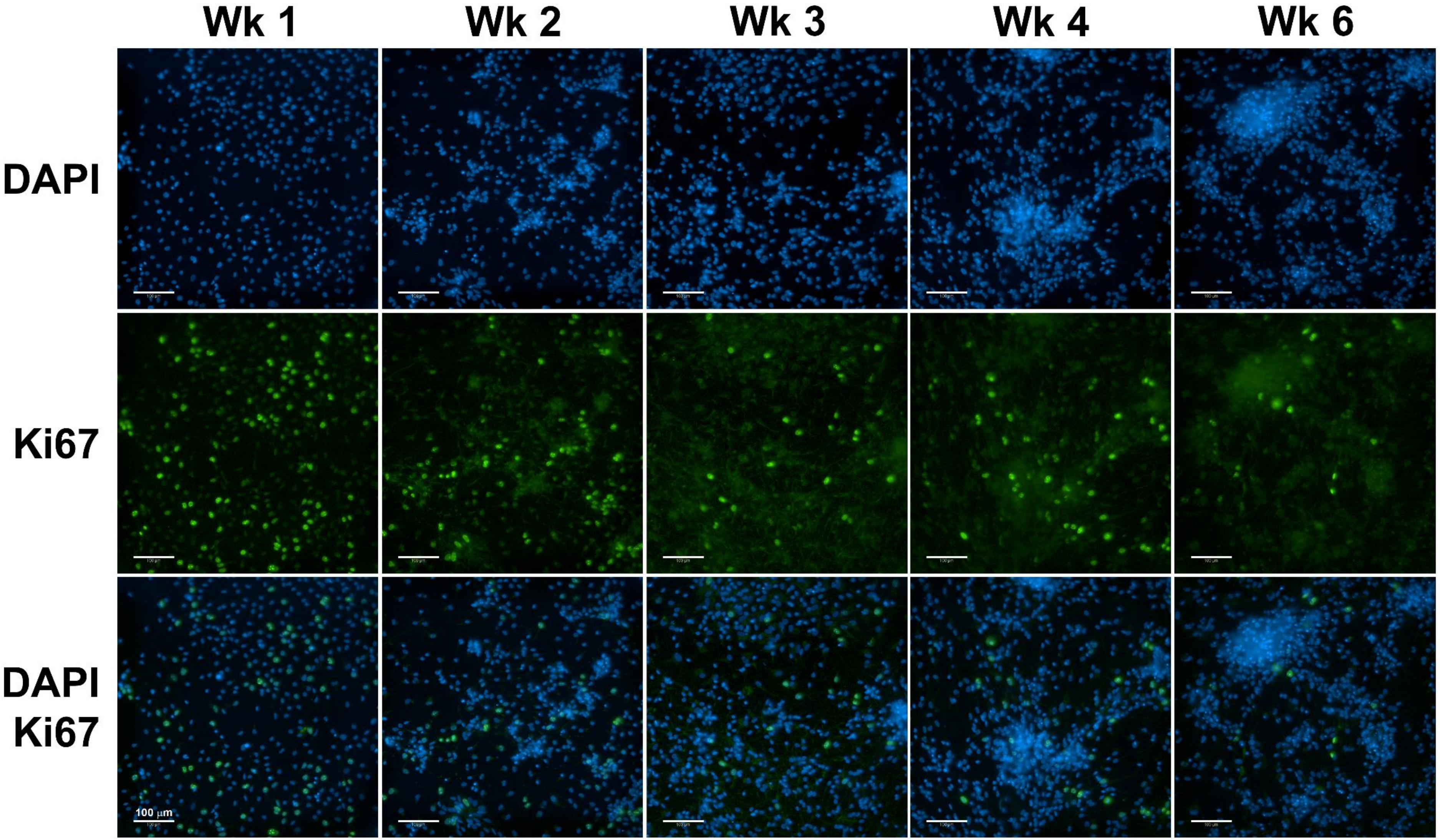


**Decrease in Ki67 gene expression during iPSC-derived neuronal differentiation and maturation under feeder layer free conditions.** Representative fields of view of Ki67 positivity in weeks 1-6 post-differentiation. Wk=week post-differentiation.

#### Supplemental Figure 4

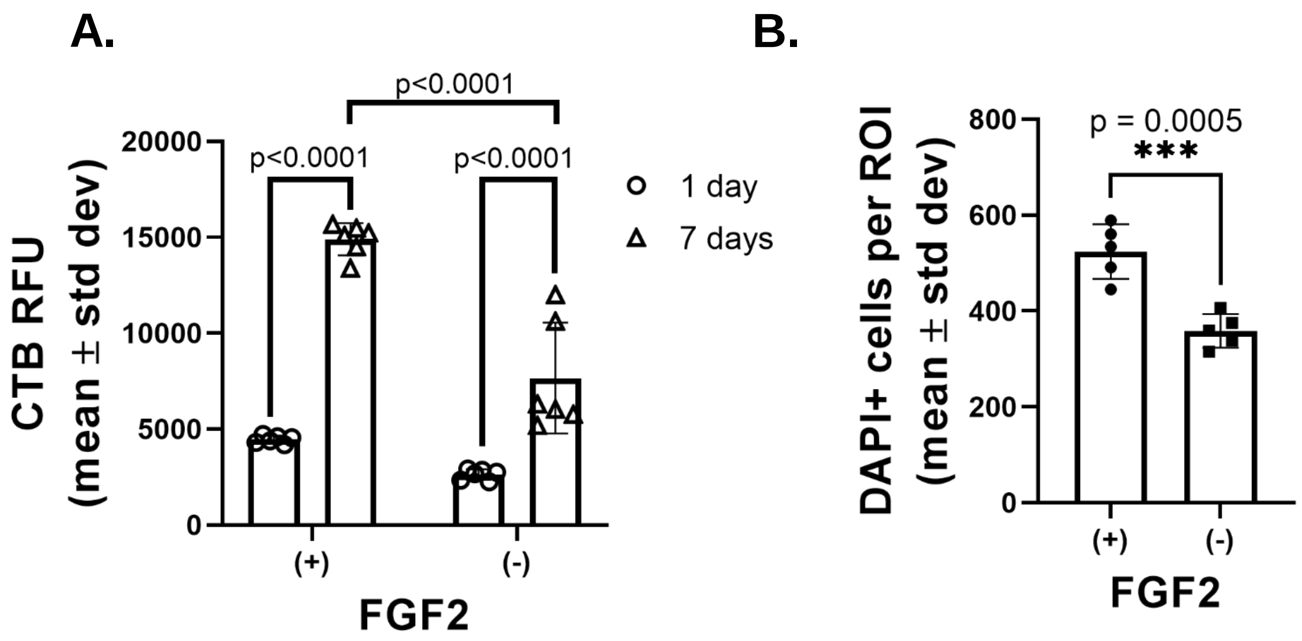


**Removal of rhFGF suppresses radial glial cell proliferation.** iPSC-derived neurogenesis is a multi-step process and initiates with the differentiation to rhFGF-responsive radial glial cells. Carryover of thermostable rhFGF from NPC cell culture contributes to the asynchronous expansion of radial glial cells and compromises cell seeding reproducibility during neuronal differentiation. Panel A, Metabolic assessments using Cell Titer Blue (CTB) at 1 and 7 days post-differentiation initiation. Panel B, DAPI+ cells after 14 days post-differentiation.

### Supplemental Figure 5
